# Genetic analysis of the rice jasmonate receptors reveals specialized function for *OsCOI2*

**DOI:** 10.1101/2022.12.12.520024

**Authors:** Hieu Trang Nguyen, Mohamad Cheaib, Marie Fournel, Maelle Rios, Pascal Gantet, Laurent Laplaze, Soazig Guyomarc’h, Michael Riemann, Thierry Heitz, Anne-Sophie Petitot, Antony Champion

## Abstract

- COI1-mediated perception of jasmonate is critical for plant development and responses to environmental stresses. Monocots such as rice have two groups of *COI* genes due to gene duplication: *OsCOI1a* and *OsCOI1b* that are functionally equivalent to the dicotyledons *COI1* on one hand and *OsCOI2* whose function remains unclear.
- In order to assess the function of *OsCOI2* and its functional redundancy with *COI1* genes, we developed a series of rice mutants in the 3 genes *OsCOI1a, OsCOI1b* and *OsCOI2* by CRISPR Cas9 and characterized their phenotype and responses to jasmonate.
- Characterization of *OsCOI2* uncovered important roles in root, leaf and flower development. In particular, we show that crown root growth inhibition by jasmonate relies on *OsCOI2* and not *OsCOI1a* or *OsCOI1b* in rice, revealing a major function for the non-canonical *OsCOI2* in jasmonate-dependent control of rice root growth.
- Collectively, these results point to a specialized function of *OsCOI2* in the regulation of plant development in rice and indicate that sub-functionalisation of jasmonate receptors has occurred in the monocot phylum.

## Introduction

Jasmonoyl-isoleucine (JA-Ile), the bioactive form of jasmonic acid (JA), regulates many physiological and developmental processes in plants such as flower and root development (Yuan & Zhang, 2015; Lakehal *et al.*, 2020). Plants from monocot and eudicot lineages accumulate JA-Ile in response to stresses, such as injury or insect attack (Lyons *et al.*, 2013; Schluttenhofer, 2020). Jasmonate perception and transcriptional regulation of JA-responsive genes relies on a conserved core protein complex, the main components of which are the F-box protein CORONATINE INSENSITIVE1 (COI1) protein, the JASMONATE ZIM DOMAIN (JAZ) transcriptional repressors and transcription factors (TFs) (Xie *et al.*, 1998; Chini *et al.*, 2007; Thines *et al.*, 2007; Yan *et al.*, 2007; Sheard *et al.*, 2010). In the absence of JA-Ile, target gene expression is repressed due to TF and JAZ protein interaction within the TOPLESS (TPL)-NOVEL INTERACTORS OF JAZ (NINJA) repressor complex (Pauwels et al 2010). Upon JA-Ile accumulation, JAZ proteins are recruited by COI1, ubiquitinated and rapidly degraded in a proteasome-dependent manner, thus freeing TFs and leading to the de-repression of JA-responsive genes and subsequent physiological responses.

In eudicot species such as Arabidopsis and tomato, JA-Ile is perceived by a single receptor encoded by *COI1* that interacts with members of the JAZ family (Xie *et al.*, 1998; Chini *et al.*, 2007; Thines *et al.*, 2007). In contrast, monocots such as rice, maize and wheat have two groups of *COI* genes due to gene duplication (An *et al.*, 2019: Schluttenhofer, 2020; Qi *et al.*, 2022). In rice, there are three *COI* genes, named *OsCOI1a, OsCOI1b*, and *OsCOI2. OsCOI1a* and *OsCOI1b* are orthologous to *AtCOI1* as shown by complementation of the loss-of-function *coi1-1* mutant in Arabidopsis (Lee *et al.*, 2013). Several studies in rice demonstrated the involvement of *OsCOI1a* and *OsCOI1b* in plant defence. Transgenic rice plants with a reduced expression of both *OsCOI1a* and *OsCOI1b* genes display a phenotype similar to gibberellic acid over-accumulating plants, manifested by elongated internodes suggesting that jasmonate prioritizes defence over growth (Yang *et al.*, 2012). In a second study, silencing of *OsCOI1a* and *OsCOIb* gene expression lead to a reduced resistance phenotype to the rice leaf-folder insect (Ye *et al.*, 2013) and to the rice stripe virus which resistance is conferred by JA signalling elements (Yang *et al.*, 2020).

In contrast, very little is known about *OsCOI2* function in rice. OsCOI2 fails to rescue the fertility and defence response defect of *coi1-1*, nor interacts with any AtJAZ co-receptor (Lee *et al.*, 2013), suggesting that OsCOI2 may have as-yet unidentified function(s) distinct from OsCOI1. Similarly, in maize (*Zea mays*), three closely related *ZmCOI1* genes could rescue *coi1-1* mutant in *Arabidopsis*, but *ZmCOI2* did not (An *et al.*, 2019). Interestingly, substitution in OsCOI2 of one amino acid predicted to be involved in JA-Ile interaction allowed it to interact with AtJAZs and rescue *coi1-1* deficiency (Lee *et al.*, 2013). Monocot and dicot lineages split about 140 to 150 million years ago (Chaw *et al.*, 2004). Phylogenetic analysis revealed that OsCOI2 shares higher similarity to COI2s from other monocots than to COI1s from monocots and eudicots, suggesting that the monocot-specific COI1-COI2 duplication and divergence occurred early in the monocot phylum (An *et al.*, 2019). This high similarity in the sequence of COI2 proteins from monocot species, and the failure of *OsCOI2* and *ZmCOI2* to complement the Arabidopsis *coi1-1* mutant raised fundamental questions about the conservation of function of *COI2* and whether it was involved in jasmonate signalling. Here genetic analysis allowed us to address in parallel the functions of OsCOI1a/b and OsCOI2, and to uncover the unique roles of OsCOI2 in jasmonate mediated developmental responses.

## Materials and Methods

### Edited rice lines

To generate the JA perception mutants *oscoi1a/b* and *oscoi2*, the CRISPR-Cas9 system was used on the background cultivar Kitaake (*Oryza sativa* L. ssp. japonica) as described in Nguyen *et al.* (2020). Briefly, the CRISPR Guides tool on Benchling platform (https://benchling.com/crispr) was used to design the gRNAs. One gRNA was designed for each gene, *OsCOI1a* and *OsCOI1b*, to obtain the *oscoi1a/b* double mutants and two different pairs of gRNAs were designed to obtain the *oscoi2* mutants. The sequences of the gRNAs were first inserted into the pUC57-sgRNA vector, then transferred by LR recombination into the binary vector pOS-Cas9 (Miao *et al.*, 2013). The pOS-Cas9-gRNAs vectors were introduced in the *Agrobacterium tumefaciens* strain EHA105 by electroporation. Rice embryogenic calli were transformed via *Agrobacterium*-mediated transformation protocols. The regenerated plants were checked for transformation, by PCR using T-DNA specific primers (HPT and Cas9, Table S1), and for mutation, by PCR using primers flanking the targeted sequences (Table S1) and sequencing (Eurofins). Edited plants were transferred to a greenhouse for multiplication. Edited lines were counter-selected from T1 generation to obtain T3 homozygous lines without T-DNA insertion, except for *oscoi2-1* for which the limited number of seeds prevented such selection.

### Plant phenotyping

Seeds were sterilized in 70% ethanol for 1 minute, then in 40% commercial bleach for 30 minutes, and were rinsed 6 times in sterilized water. Disinfected seeds were then germinated on half-strength Murashige & Skoog (½MS) medium including Gamborg B5 vitamins (Duchefa Biochemie BV, Haarlem, the Netherlands) supplied with 0.7% plant agar (Duchefa) for 6 days. Plantlets of uniform size were transferred into soil and grown in a greenhouse under a 16h-day/8h-night photoperiod, at 28°C/24°C with a 75% relative humidity. Tiller and internode lengths were measured at the mature stage on the 3 highest tillers from 18 plants. Fertility rate was measured on the same tillers as the ratio between the number of fertile spikelets and the total number of spikelets. Leaf necrotic lesions and adventitious roots were observed under an Axiozoom microscope (Zeiss). Spikelet and anther morphology were observed under a stereo microscope (Nikon SMZ1500).

### Hormone treatments

Seeds were sterilized and sown as described above. Two days after germination, seedlings with both coleoptile and primary root were transferred into glass tubes containing 20 ml of ½MS including Gamborg B5 vitamins supplied with 0.2% phytagel (Sigma) and grown in a growth chamber at 26°C under a 12h-day/12h-night photoperiod with a 70% relative humidity. For phenotyping, 5 μM JA (Sigma), 0.5 μM COR (Sigma) both dissolved in DMSO, or DMSO only (control plants) were added to the media just before pouring. Eight days after transplanting, length of the crown roots was measured for each plant. For each line, 20-24 plants were used except for the *oscoi2-1* line for which very few seeds were available. Percentages of root growth inhibition were determined by randomly comparing the length values of the treated plants to those of control plants.

For gene expression analyses, plants were transferred 4 days after transplantation from the tubes into liquid media containing either 5 μM JA or DMSO for control plants. Crown root tips (around 1 cm) from 20-24 plants were collected 6 hours after and immediately frozen in liquid nitrogen. Five independent biological samples were collected and analysed.

### Gene expression analyses

Total RNAs were extracted from rice root samples using the RNeasy Plant mini kit (Qiagen, France), with addition of an on-column DNase I digestion. First-strand cDNAs were synthesized from 1 μg of total RNA in 20 μl final volume using an oligo-dT(18)-MN primer (Eurogentec, France) and the Omniscript RT kit (Qiagen). Specific primers were designed from *O. sativa* Nipponbare sequences using Primer3Plus (https://primer3plus.com/) (Table S1) and checked for specificity on *O. sativa* Kitaake sequences. Quantitative-PCR assays were performed on cDNAs samples (diluted 1/50e) in an Mx30005P thermal cycler (Stratagene, USA) using the Brilliant III Ultra-fast SYBR® Green QPCR Master mix with low ROX (Agilent, Santa Clara, CA, USA). Amplifications were performed in duplicate from the five biological samples. The *EXP* gene (LOC_Os06g11070) was used as a reference gene to normalize data. Relative gene expression levels were calculated using the 2^-ΔΔCt^ method.

### Statistical Analyses

Experimental data were analysed using GraphPad Prism (version 9.3.0). Values were considered statistically significant when *p* ≤ 0.05.

## Results

### *OsCOI2* regulates plant development and its function is not redundant with *OsCOI1a* and *OsCOI1b*

To investigate *COI2* function in monocots, CRISPR-Cas9 was used to generate *oscoi2* mutant alleles in rice. Two CRISPR-Cas9 constructs targeting the first (target 1 and 2), and the second exon (target 3 and 4) of *OsCOI2*, respectively, were used to generate edited lines in the rice cultivar Kitaake (Fig. 1a). Three homozygous mutant lines, named *oscoi2-1, oscoi2-2* and *oscoi2-3*, were identified and further characterized. The alleles *oscoi2-1* and *oscoi2-2* include short nucleotide insertions and deletions in the first and second exon leading to a translational frame shift and premature stop codon truncating 467 amino acids and 372 amino acids in the predicted proteins, respectively. The third allele *oscoi2-3* displays fragments deleted in exon 2 eliminating 18 amino acids in the LRR domain, which can affect the protein stability (Fig. 1b) (Yan *et al.*, 2009). In order to assess functional redundancy between OsCOI2 and OsCOI1, we also generated two double *oscoi1a oscoi1b* mutant lines (thereafter named *oscoi1a-1/b-1* and *oscoi1a-1/b-2*) by CRISPR-Cas9 (Fig. S1).

**Figure 1.**
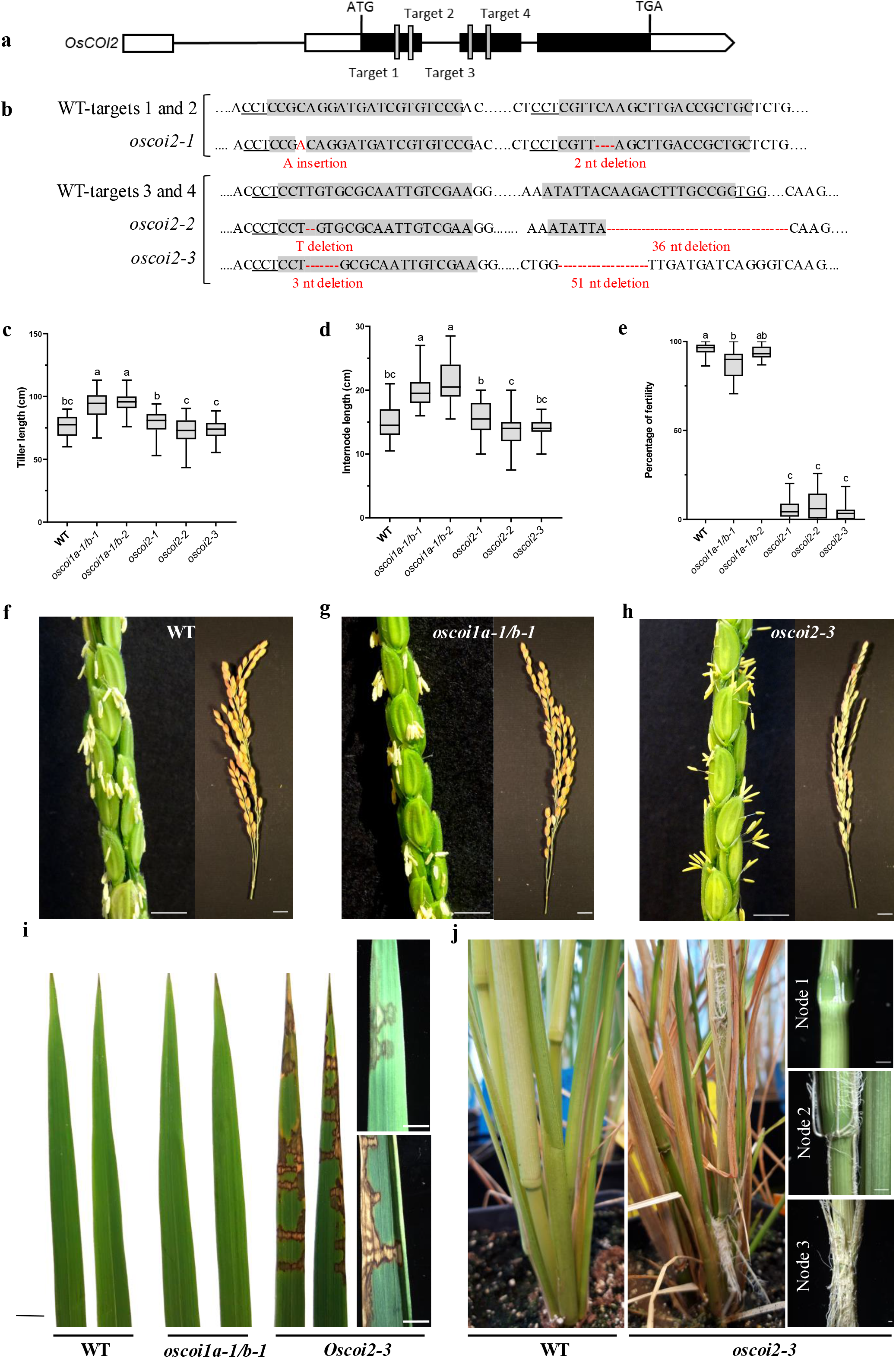
*OsCOI2* regulates vegetative and reproductive developmental programmes in rice distinct from *OsCOI1a/b*. **a**, *OsCOI2* gene structure and CRISPR-Cas9 target sites. **b**, insertion and deletion sites of three allelic mutations (*oscoi2-1, oscoi2-2* and *oscoi2-3*) generated by two different pairs of gRNA. **c**, quantification of tiller length and **d**, quantification of internode length. Letters indicate significant differences between lines (45<n<54, One way-ANOVA with Tuckey’s multiple comparisons test, *p*<0.05). **e**, Percentage of fertile spikelet. Letters indicate significant differences between lines (43<n<53, Kruskall-Wallis multiple comparisons test, *p*<0.05). In the boxplots, whiskers denote minimum/maximum values, the box defines the interquartile range and the centre line represents the median. (c, d, e). **f**-**h** images of WT (**f**), *oscoi1a-1/b-1* (**g**) and *oscoi2-3* (**h**) panicles. **i**, flag leaves. **j**, nodes. (**f**-**j**) Scale bar = 1 cm.

Phenotypically, the *oscoi1a-1/b-1*, *oscoi1a-1/b-2* lines exhibited increased plant height in comparison with *oscoi2* and wild type (WT) plants (Fig. 1c, Fig. S2 and S3). In addition, internode length was increased in the *oscoi1a-1/b-1* and *oscoi1a-1/b-2* compared to WT and *oscoi2* mutant lines (Fig. 1d, Fig. S2). These *oscoi1a/b* phenotypes are similar to those reported for the rice *coi1-RNAi* lines which suppress both *OsCOI1a* and *OsCOI1b* expression (Yang *et al.*, 2012). Thus, we used these new *oscoi1a-1/b-1* and *oscoi2* knockout lines to compare OsCOI1a/b and OsCOI2 regulation of plant development and JA signalling in rice.

The most prominent phenotype observed in *Arabidopsis coi1-1* and tomato *jai1-1* JA receptor mutants is sterility (Xie *et al.*, 1998; Li *et al.*, 2004), but earlier studies reported only a weak reduction in fertility in *oscoi1-RNAi* and *oscoi1b-T-DNA* rice mutant lines (Yang *et al.*, 2012; Lee *et al.*, 2015). Consistent with these reports, seed-setting in our *oscoi1a/b* double mutant lines was close to WT level (Fig. 1e, 1f and 1g). In contrast, all 3 of our independent *oscoi2* mutant alleles conferred reduced seed setting from T0 to T3 generations (Fig. 1e and 1h). None of the *oscoi1a/b*, nor *oscoi2* mutant lines displayed any abnormality in floret architecture (Fig. S4); however, most of the *oscoi2* anthers did not dehisce (Fig. 1f, 1g and 1h and Fig. S4). Hence, *OsCOI2*, rather than *OsCOI1a* or *OsCOI1b*, plays an important role during rice dehiscence stage of reproductive development. Importantly, even though *OsCOI2* does not restore fertility in the *Arabidopsis coi1-1* mutant background (REF), our results indicate that *OsCOI2* is likely the major regulator of jasmonate-dependent fertility in rice.

During the vegetative to reproductive transition, *oscoi2* mutant lines exhibited three distinct phenotypes not observed in WT or in *oscoi1a/b* plants. First, all three *oscoi2* mutant alleles showed spontaneous lesions on flag leaves and the last leaf mimicking disease symptoms, despite plants not being infected (Fig. 1.i). Second, when the main *oscoi2* panicle started to flower, above ground adventitious roots emerged from node 1 to node 3 that resemble adventitious roots induced by flooding in deep-water rice (Lorbiecke & Sauter, 1999). Third, 25 to 35% of the plants showed leaf rolling followed by tiller senescence and eventually plant death without imposing any water stress (Fig. S5). Collectively, these data reveal that *OsCOI2* regulates plant development and that its function is not redundant with *OsCOI1a* and *OsCOI1b* jasmonate receptors.

### *OsCOI2* is required for JA perception in rice crown roots

We next tested the hypothesis that *OsCOI2* participates in JA signalling in rice. Inhibition of seedling root growth has been extensively used as a bioassay to identify jasmonate responsive mutants in plants (Xie *et al.*, 1998; Li *et al.*, 2004; Cao *et al.*, 2021). Under our root growth conditions, JA at 5 μM inhibits similarly root growth of WT plants and *oscoi1a/b* double mutant lines (Fig. 2a and 2b). In contrast, crown root growth was significantly less sensitive to exogenous JA treatment in *oscoi2-1, oscoi2-2*, and *oscoi2-3* genetic backgrounds (Fig. 2a and 2b and Fig. S6). Similarly, coronatine (COR), a bacterial mimic of JA-Ile, strongly inhibited root growth of WT and *oscoi1a/b* lines but had a significantly weaker effect on root growth of *oscoi2* mutant alleles (Fig. 2a and 2b).

**Figure 2.**
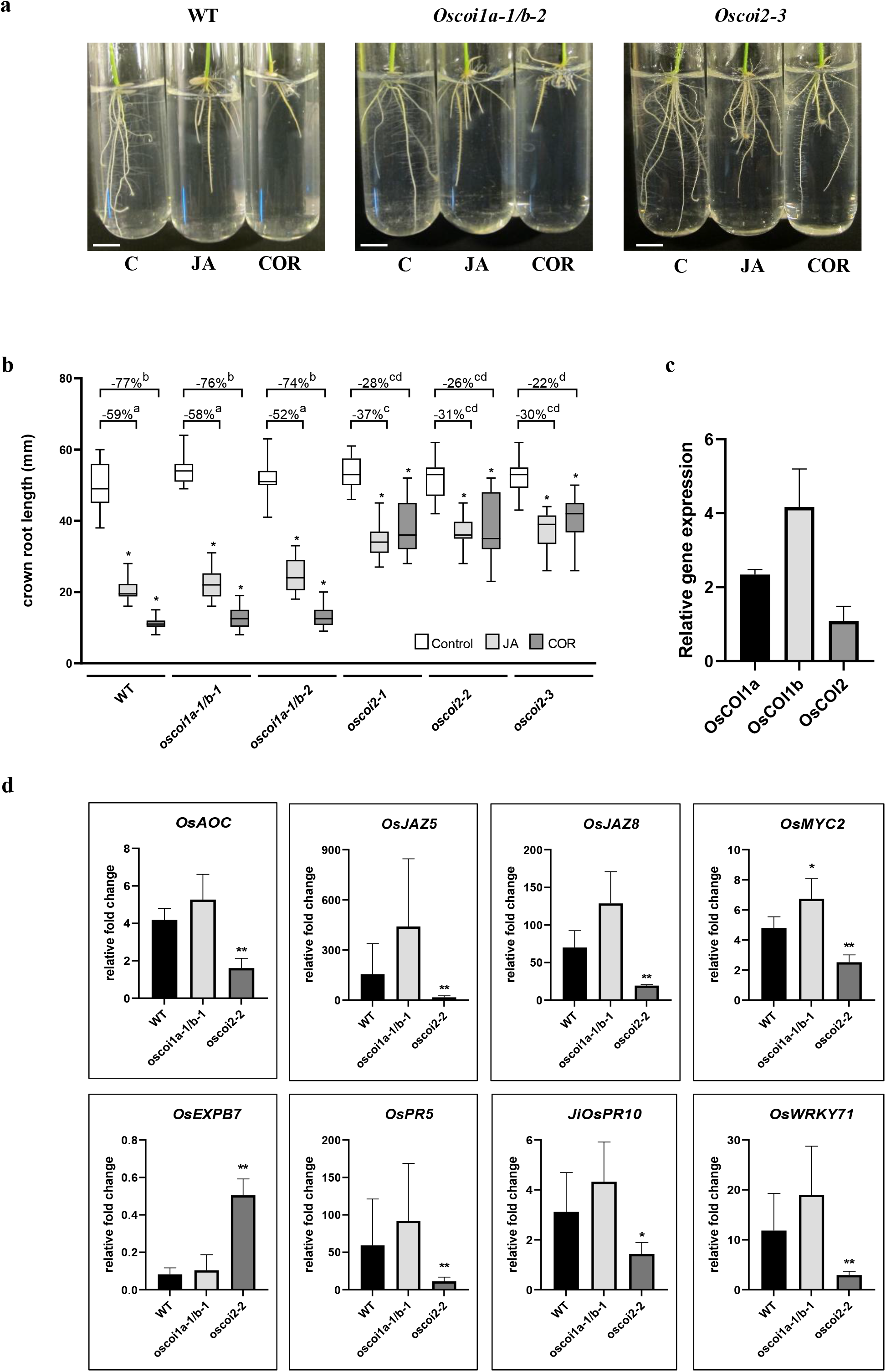
*OsCOI2* regulates jasmonate response in crown roots. **a**, Phenotype of the (WT) and two allelic mutations (*oscoi1a-1/b-2* and *oscoi2-3*) of roots in response to jasmonic acid (JA, 5 μM), coronatine (COR, 0.5 μM) and mock control (C, DMSO). Scale bars = 1 cm. **b**, effect of JA and COR compared to the mock treatment (DMSO) on crown root length in WT *oscoi1a/b* (*oscoi1a-1/b-1* and *oscoi1a-1/b-2*) and *OsCOI2* (*oscoi2-1, oscoi2-2* and *oscoi2-3*) mutant lines. In the boxplots, whiskers denote minimum/maximum values, the box defines the interquartile range and the centre line represents the median. Asterisks indicate significant differences between treated and control plants (20<n<24 except for *oscoi2-1* where 7<n<21, One way-ANOVA with Bonferroni’s multiple comparisons test, *p*<0.05). The percentages of crown root growth inhibition by JA and COR treatments are indicated above the respective plots from the WT, *oscoi1a-1/b-1*, *oscoi1a-1/b-2, oscoi2-1, oscoi2-2* and *oscoi2-3* mutants. Letters indicate significant differences between lines and treatments (21<n<24, One way-ANOVA with Tuckey’s multiple comparisons test, *p*<0.05). **c.** expression of *OsCOI1a, OsCOI1b* and *OsCOI2* genes in crown root tips from wild-type plants. Data are presented as the means +/- SD from five biological replicates. **d**, regulatory effects of JA on the transcription of jasmonate responsive genes and defence genes. Data are presented as means +/- SD from five biological replicates. Asterisks indicate significant differences between the WT and the double *oscoi1a-1/b-1* mutant or the *oscoi2-2* mutant (Mann-Whitney tests, * *p* <0.05, \*\**p*<0.01).

To explore the molecular mechanisms associated with inhibition of crown root growth by jasmonate, we focused on crown root tips, where cell proliferation and elongation occur. All three JA receptor genes *OsCOI1a, OsCOI1b and OSCOI2* were expressed in crown root tips (Fig. 2c). Induction of jasmonate responsive genes related to JA biosynthesis (*OsAOC*) and signalling (*OsJAZ5, OsJAZ8* and *OsMYC2*) by exogenous JA application was compromised in *oscoi2-2* crown roots relative to wild type, but not in *oscoi1a-1/b-1* mutant background (Fig. 2d, top panels). Consistent with the root growth phenotype, expression of the cell elongation-related gene *OsEXPB7* was significantly less repressed by JA treatment in *oscoi2-2* crown roots than in wild type and *oscoi1a-1/b-1* mutant (Fig. 2d) (Zou *et al.*, 2021). Furthermore, genes associated with defence responses (*OsPR5, OsPR10* and *OsWRKY71*) were significantly less induced by JA treatment in *oscoi2-2* than in WT and *oscoi1a-1/b-1*. Hence, expression profiling revealed that *OsCOI2*, and not *OsCOI1a/b*, is required for JA-dependent signalling in rice crown roots.

## Discussion

Here, we report that a monocot-specific member of the COI family *OsCOI2* mediates the regulation of a range of jasmonate-dependent vegetative and reproductive developmental processes. Significantly, loss of *OsCOI2* function (and not *OsCOI1a* and *OsCOI1b*) was shown to strongly alter male fertility in rice, a phenotype that is *AtCOI1*-dependent in Arabidopsis. In rice, jasmonate is not only necessary for stamen fertility, but also required for floret development (Cai *et al.*, 2014; Nguyen *et al.*, 2019). Loss-of-function mutations or misexpression in key genes in the jasmonate biosynthetic or signalling pathways cause floral morphological alterations such as longer sterile lemmas and extra glume-like organs (Cai *et al.*, 2014; Nguyen *et al.*, 2019; Cao *et al.*, 2021). Here, we show that *oscoi2* anthers did not dehisce properly, a phenotype also observed in the *osjar1* mutant impaired in JA-Ile biosynthesis (Xiao *et al.*, 2014). Furthermore, previous reports showed that *OsCOI2* does not restore fertility and JA signal transduction in the Arabidopsis *coi1* mutant, unless the *OsCOI2* His-391 was substituted with Tyr-391 (Lee *et al.*, 2013). Collectively, these data point to a specialized function of COI2 in the regulation of plant development in the Monocot phylum.

*COI1* was shown to be the receptor of the jasmonate signal in Arabidopsis and in rice. We investigated the possible role of *OsCOI2* in jasmonate signalling using the JA-mediated repression of crown root growth. Surprisingly, we discovered that inhibition of crown root growth by JA relies on *OsCOI2* and not *OsCOI1a* or *OsCOI1b* in rice. Consistently, expression profiling revealed that *OsCOI2* is specifically required for JA-dependent signalling in crown roots. This suggests that sub-functionalisation of JA receptors has occurred in the monocot lineage.

Besides its role in root growth inhibition, jasmonate is a potential regulator of adventitious root initiation. For example, JA treatment inhibited adventitious root formation in Arabidopsis hypocotyl (Gutierrez *et al.*, 2012). Accordingly, *atcoi1-1* mutant produced more adventitious root compared to the wild type indicating a negative role of JA in this developmental process (Gutierrez *et al.*, 2012). In rice, adventitious root emergence is inducible by flooding and is tightly regulated by hormones such as ethylene and auxin (Mergemann & Sauter, 2000; Lin & Sauter, 2019). Constitutive formation of adventitious roots from *oscoi2* nodes suggests that this non-canonical jasmonate receptor could be a central regulator of rhizotaxis plasticity in rice.

Specific functions of *OsCOI2* could derive from (i) its spatial expression pattern being different from *OsCOI1a/b*, and/or (ii) distinct protein-protein interactions involving a subset of OsJAZs, and/or (iii) the perception of distinct jasmonate forms. Recently, the liverwort *Marchantia polymorpha* MpCOI1 receptor was shown to interact with distinct hormone forms that the eudicot Arabidopsis COI1 receptor did not bind (Monte *et al.*, 2018). Specifically, instead of perceiving JA-Ile, MpCOI1 binds the JA precursor dn-OPDA as an ancestral ligand to initiate jasmonate signalling (Monte *et al.*, 2018). In rice, despite of JA-Ile functions being established, OPDA- and distinct jasmonic acid-amino acid conjugates have been shown recently to be involved in defence response and could be additional ligands of OsCOI proteins Shinya *et al.*, 2021; Fu *et al.*, 2022.

In summary, we report that COI jasmonate receptor sub-functionalisation has occurred in rice, consistent with the hypothesis that different bioactive forms of jasmonate could modulate distinct responses to developmental cues and environmental stresses via *OsCOI1a/b* and *OsCOI2* perception. Our collection of *oscoi* mutant alleles offers a unique genetic resource for future dissection of the function of jasmonate receptor proteins in monocots.

## Acknowledgements

This work was supported by the Consultative Group for International Agricultural Research Program on rice-agrifood systems (CRP-RICE, 2017–2022). HTN is funded by a PhD fellowship from the French Embassy in Vietnam. We thank B.K. Pandey and M. Bennett for advice while drafting the manuscript. We thank E. Guillon and P. Serin for technical assistance with the IRD glasshouse work.

## Author Contributions

H.T.N., A.S.P., and A.C. designed the experiments. H.T.N., M.C., M.F., M.R., A.S.P, and A.C. performed the experiments, and H.T.N., A.S.P., M.R., T.H., S.G., P.G., L.L., and A.C. analysed the data. H.T.N., A.S.P., and A.C. wrote the manuscript with contributions from all the other authors.

## Supporting information

**Figure S1.** Schematic representation of the *oscoi1ab* double mutant lines.

**Figure S2.** Images of wild type, *oscoi1a-1/b-1*, *oscoi1a-1/b-2*, *oscoi2-2* and *oscoi2-3* show panicule phenotypes.

**Figure S3.** Quantification of stem length of the WT and *oscoi* mutant plantlets.

**Figure S4.** Comparison of anther dehiscence between the wild-type and the *oscoi* mutants.

**Figure S5.** Image of wild-type and *oscoi2-3* mutant plants showing mortality at reproductive stage.

**Figure S6.** Crown root growth inhibition assay by JA application on WT and T2 *oscoi* mutants.

**Table S1.** List of primers used for qPCR and plant genotyping.

## References

An L, Ahmad RM, Ren H, Qin J, Yan Y. 2019. Jasmonate Signal Receptor Gene Family *ZmCOIs* Restore Male Fertility and Defense Response of *Arabidopsis* mutant *coi1-1*. Journal of Plant Growth Regulation 38: 479–493.

Cai Q, Yuan Z, Chen M, Yin C, Luo Z, Zhao X, Liang W, Hu J, Zhang D. 2014. Jasmonic acid regulates spikelet development in rice. Nature Communications 5: 3476–3489.

Cao L, Tian J, Liu Y, Chen X, Li S, Persson S, Lu D, Chen M, Luo Z, Zhang D et al. 2021. Ectopic expression of *OsJAZ6*, which interacts with OsJAZ1, alters JA signaling and spikelet development in rice. Plant Journal 108: 1083–1096.

Chaw SM, Chang CC, Chen HL, Li WH. 2004. Dating the monocot-dicot divergence and the origin of core eudicots using whole chloroplast genomes. Journal of Molecular Evolution 58: 424–441.

Chini A, Fonseca S, Fernández G, Adie B, Chico JM, Lorenzo O, García-Casado G, López-Vidriero I, Lozano FM, Ponce MR et al. 2007. The JAZ family of repressors is the missing link in jasmonate signalling. Nature 448: 666–671.

Fu W, Jin G, Jiménez-Alemán GH, Wang X, Song J, Li S, Lou Y, Li R. 2022. The jasmonic acid-amino acid conjugates JA-Val and JA-Leu are involved in rice resistance to herbivores. Plant Cell & Environment 45: 262–272.

Gutierrez L, Mongelard G, Floková K, Pacurar DI, Novák O, Staswick P, Kowalczyk M, Pacurar M, Demailly H, Geiss G, Bellini C. 2012. Auxin controls Arabidopsis adventitious root initiation by regulating jasmonic acid homeostasis. Plant Cell 24:2515–2527.

Lakehal A, Ranjan A, Bellini C. 2020. Multiple Roles of Jasmonates in Shaping Rhizotaxis: Emerging Integrators. Methods in Molecular Biology 2085: 3–22.

Lee HY, Seo JS, Cho JH, Jung H, Kim JK, Lee JS, Rhee S, Do Choi Y. 2013. *Oryza sativa* COI homologues restore jasmonate signal transduction in *Arabidopsis coi1-1* mutants. PLoS One 8: e52802.

Lee SH, Sakuraba Y, Lee T, Kim KW, An G, Lee HY, Paek NC. 2015. Mutation of *Oryza sativa CORONATINE INSENSITIVE 1b* (*OsCOI1b*) delays leaf senescence. Journal of Integrative Plant Biology 57: 562–576.

Li L, Zhao Y, McCaig BC, Wingerd BA, Wang J, Whalon ME, Pichersky E, Howe GA. 2004. The tomato homolog of CORONATINE-INSENSITIVE1 is required for the maternal control of seed maturation, jasmonate-signaled defense responses, and glandular trichome development. The Plant Cell 16: 126–143.

Lin C, Sauter M. 2019. Polar Auxin Transport Determines Adventitious Root Emergence and Growth in Rice. Frontiers in Plant Science 10:444.

Lorbiecke R, Sauter M. 1999. Adventitious root growth and cell-cycle induction in deepwater rice. Plant Physiology 119: 21–30.

Lyons R, Manners JM, Kazan K. 2013. Jasmonate biosynthesis and signaling in monocots: a comparative overview. Plant Cell Reports 32: 815–827.

Mergemann H, Sauter M. 2000. Ethylene induces epidermal cell death at the site of adventitious root emergence in rice. Plant Physiology 124: 609–614.

Miao J, Guo D, Zhang J, Huang Q, Qin G, Zhang X, Wan J, Gu H, Qu LJ. 2013. Targeted mutagenesis in rice using CRISPR-Cas system. Cell Research 23: 1233–1236.

Monte I, Ishida S, Zamarreño AM, Hamberg M, Franco-Zorrilla JM, García-Casado G, Gouhier-Darimont C, Reymond P, Takahashi K, García-Mina JM et al. 2018. Ligand-receptor co-evolution shaped the jasmonate pathway in land plants. Nature Chemical Biology 14: 480–488.

Nguyen TH, Thi Mai To H, Lebrun M, Bellafiore S, Champion A. 2019. Jasmonates-the Master Regulator of Rice Development, Adaptation and Defense. Plants. 8: 339.

Nguyen TH, Mai HTT, Moukouanga D, Lebrun M, Bellafiore S, Champion A. 2020. CRISPR/Cas9-Mediated Gene Editing of the Jasmonate Biosynthesis *OsAOC* Gene in Rice. Methods in Molecular Biology 2085: 199–209.

Pauwels L, Barbero GF, Geerinck J, Tilleman S, Grunewald W, Pérez AC, Chico JM, Bossche RV, Sewell J, Gil E, García-Casado G, Witters E, Inzé D, Long JA, De Jaeger G, Solano R, Goossens A. 2010. NINJA connects the co-repressor TOPLESS to jasmonate signalling. Nature 464: 788–791.

Qi X, Guo S, Wang D, Zhong Y, Chen M, Chen C, Cheng D, Liu Z, An T, Li J et al. 2022. *ZmCOI2a* and *ZmCOI2b* redundantly regulate anther dehiscence and gametophytic male fertility in maize. Plant Journal, in press.

Schluttenhofer C. 2020. Origin and evolution of jasmonate signaling. Plant Science 298: 110542.

Sheard LB, Tan X, Mao H, Withers J, Ben-Nissan G, Hinds TR, Kobayashi Y, Hsu FF, Sharon M, Browse J et al. 2010. Jasmonate perception by inositol-phosphate-potentiated COI1-JAZ co-receptor. Nature 468: 400–405.

Shinya T, Miyamoto K, Uchida K, Hojo Y, Yumoto E, Okada K, Yamane H, Galis I. 2021. Chitooligosaccharide elicitor and oxylipins synergistically elevate phytoalexin production in rice. Plant Molecular Biology doi: 10.1007/s11103-021-01217-w.

Thines B, Katsir L, Melotto M, Niu Y, Mandaokar A, Liu G, Nomura K, He SY, Howe GA, Browse J. 2007. JAZ repressor proteins are targets of the SCF(COI1) complex during jasmonate signalling. Nature 448: 661–665.

Xiao Y, Chen Y, Charnikhova T, Mulder PP, Heijmans J, Hoogenboom A, Agalou A, Michel C, Morel JB, Dreni L et al. 2014. *OsJAR1* is required for JA-regulated floret opening and anther dehiscence in rice. Plant Molecular Biology 86: 19–33.

Xie DX, Feys BF, James S, Nieto-Rostro M, Turner JG. 1998. *COI1:* an *Arabidopsis* gene required for jasmonate-regulated defense and fertility. Science 280: 1091–1094.

Yan J, Zhang C, Gu M, Bai Z, Zhang W, Qi T, Cheng Z, Peng W, Luo H, Nan F et al. 2009. The *Arabidopsis* CORONATINE INSENSITIVE1 protein is a jasmonate receptor. Plant Cell 21: 2220–2236.

Yan Y, Stolz S, Chételat A, Reymond P, Pagni M, Dubugnon L, Farmer EE. 2007. A downstream mediator in the growth repression limb of the jasmonate pathway. Plant Cell 19: 2470–2483.

Yang DL, Yao J, Mei CS, Tong XH, Zeng LJ, Li Q, Xiao LT, Sun TP, Li J, Deng XW et al. 2012. Plant hormone jasmonate prioritizes defense over growth by interfering with gibberellin signaling cascade. Proceedings of the National Academy of Sciences of the United States of America 109: 1192–1200.

Yang Z, Huang Y, Yang J, Yao S, Zhao K, Wang D, Qin Q, Bian Z, Li Y, Lan Y et al. 2020. Jasmonate Signaling Enhances RNA Silencing and Antiviral Defense in Rice. Cell Host & Microbe 28: 89–103.

Ye M, Song Y, Long J, Wang R, Baerson SR, Pan Z, Zhu-Salzman K, Xie J, Cai K, Luo S et al. 2013. Priming of jasmonate-mediated antiherbivore defense responses in rice by silicon. Proceedings of the National Academy of Sciences of the United States of America 110: 3631–3639.

Yuan Z, Zhang D. 2015. Roles of jasmonate signalling in plant inflorescence and flower development. Current Opinion in Plant Biology 27: 44–51.

Zou X, Liu L, Hu Z, Wang X, Zhu Y, Zhang J, Li X, Kang Z, Lin Y, Yin C. 2021. Salt-induced inhibition of rice seminal root growth is mediated by ethylene-jasmonate interaction. Journal of Experimental Botany 72: 5656–5672.

