## Supplementary material for "Genetic analysis of the rice jasmonate receptors reveals specialized function for *OsCOI2*": Figure S1

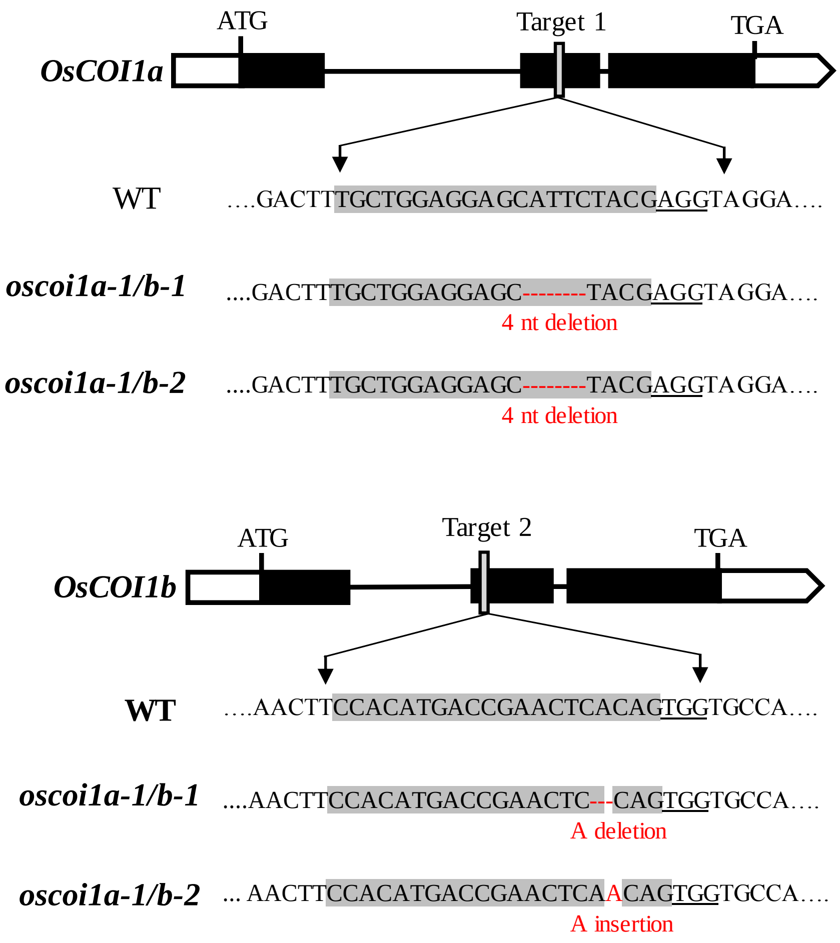


**Figure S1. Schematic representation of the *oscoi1ab* double mutant lines.**
*OsCOI1a and OsCOI1b* gene structures and the CRISPR-Cas9 target sites are shown. The insertion and deletion sites of two allelic mutations (*oscoi1a-1/b-1*, *oscoi1a-1/b-2*) generated by two gRNAs are shown in comparison with the wild-type (WT) sequence.

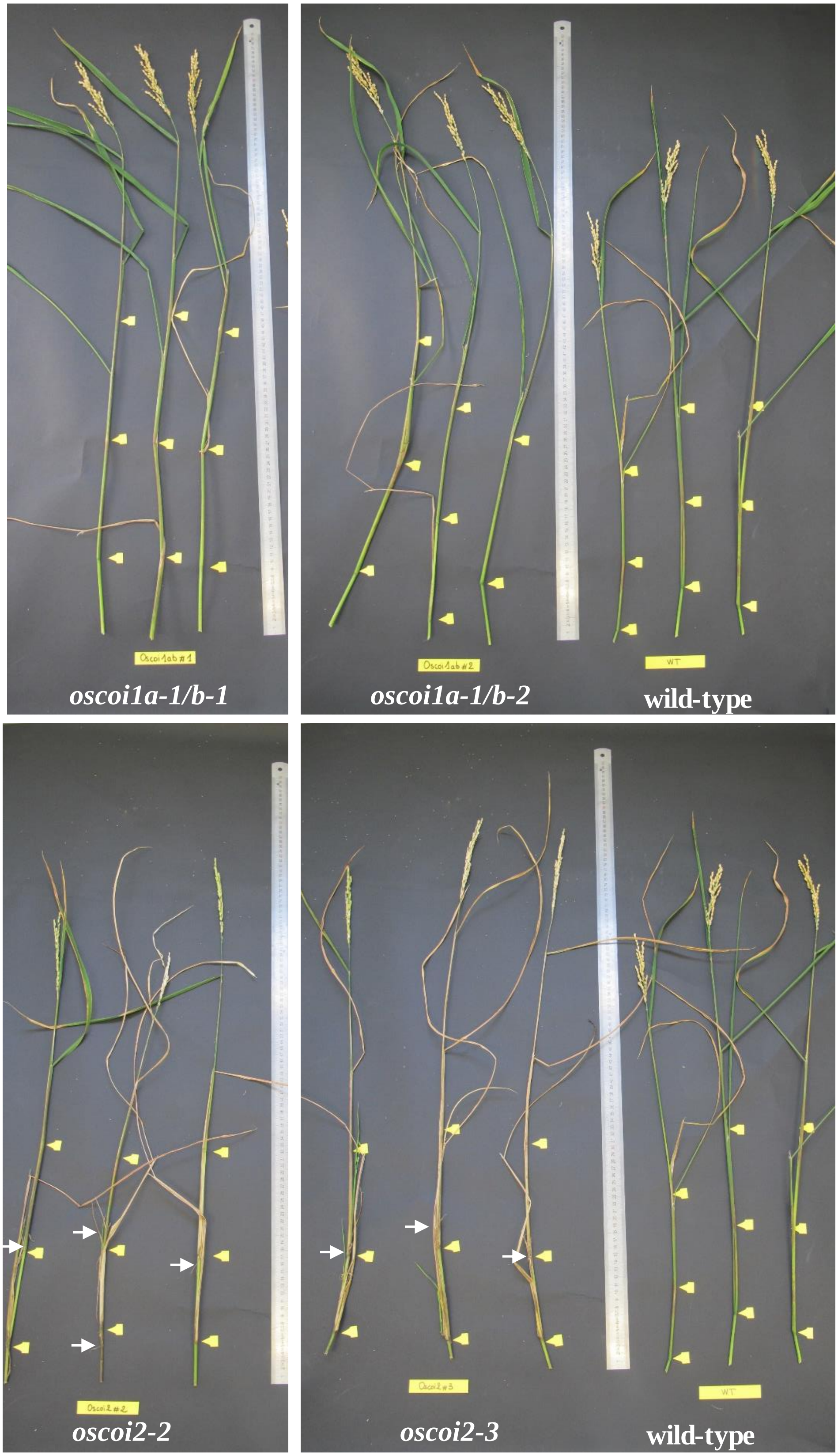


**Figure S2. Images of wild type, *oscoi1a-1/b-1*, *oscoi1a-1/b-2*, *oscoi2-2* and *oscoi2-3* show panicule phenotypes.**
Yellow arrows show nodes 1, 2 and 3. White arrows indicate adventitious roots from nodes.

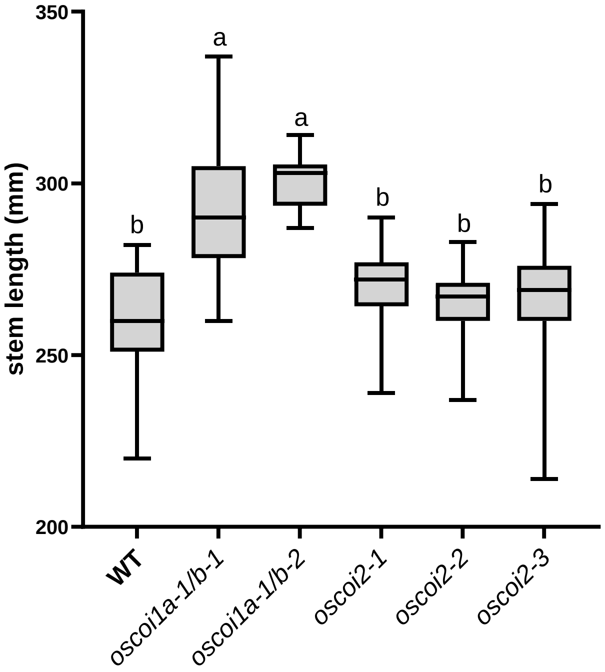


**Figure S3. Quantification of stem length of the WT and *oscoi* mutant plantlets.**

Quantification of stem length of the WT and *oscoi* mutants plantlets. Letters indicate significant differences between lines (21<n<24, One way-ANOVA with Tuckey's multiple comparisons test, $p<0.05$). In the boxplots, whiskers denote minimum/maximum values, the box defines the interquartile range and the centre line represents the median.

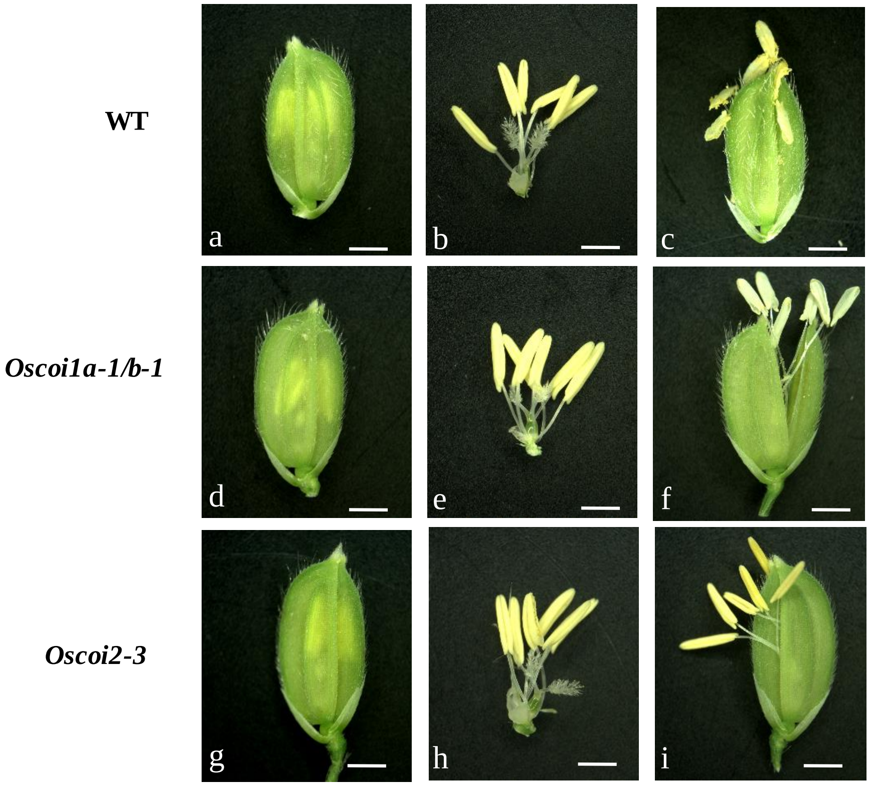


**Figure S4. Comparison of anther dehiscence between the wild-type and the *oscoi* mutants.**
Close spikelet before anthesis (a, b ,d, e, g and h) and after widely opened spikelet (c, f, i).

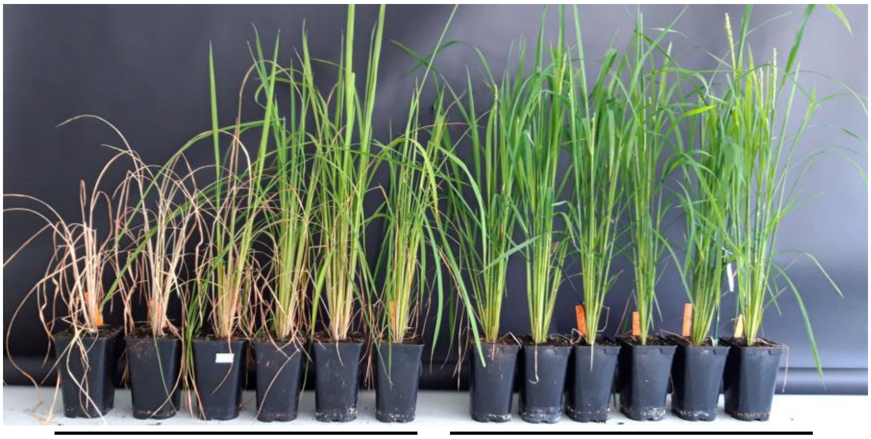


**Figure S5. Image of wild-type and *oscoi2-3* mutant plants showing mortality at reproductive stage.**

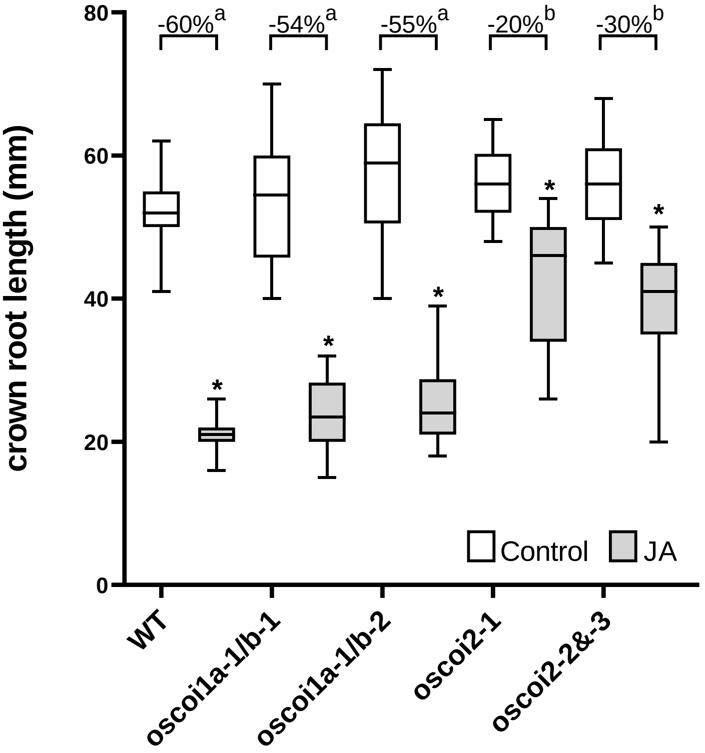


**Figure S6. Crown root growth inhibition assay by JA application on WT and T2 *oscoi* mutants.** Effect of JA on crown root length of the WT, two allelic mutations of the double mutant *oscoi1a*/*oscoi1b* (*oscoi1a-1/b-1* and *oscoi1a-1/b-2*) and two mutants of *OsCOI2* (*oscoi2-1, oscoi2-2&*-3 heterozygous line). In the boxplots, whiskers denote minimum/maximum values, the box defines the interquartile range and the center line represents the median. Asterisks indicate significant differences between treated and control plants (One way-ANOVA with Bonferroni's multiple comparisons test, $p<0.05$).The plot values are the means of the crown roots. Percentage of crown root growth inhibition by JA from the WT, *oscoi1a-1/b-1*, *oscoi1a-1/b-2, oscoi2-1* and *oscoi2-2&*-3 heterozygous line. Letters (a, b) indicate significant differences between lines (One way-ANOVA with Tuckey's multiple comparisons test, $p<0.05$)

| Gene name | Locus or Target | Forward primer | Reverse primer | Amplicon size | Use |
|---|---|---|---|---|---|
| *OsCOI1a* | LOC_Os01g63420 | GATGCCCTCCCTGAGATACA | CCACACAGGGTTCTCCATCT | 158 bp | qPCR |
| *OsCOI1b* | LOC_Os05g37690 | CAGGCCTTGGCTATATTGGA | CAGGGAAGGCATACTCCGTA | 188 bp | qPCR |
| *OsCOI2* | LOC_Os03g15880 | ATGGGTGCAAGGATACAAGG | TGCAAGAATCTGTGCCTGAC | 133 bp | qPCR |
| *OsAOC* | LOC_Os03g32314 | AAGAGGAATCGAGGACAAGATATTTG | AAGCCTCTTCTTGTTCGGATCA | 75 bp | qPCR |
| *OsJAZ5* | LOC_Os04g32480 | CGAGGCAACTAAAGCAAAAGGA | TGAGTGGCTCTTTGGCAAAATT | 71 bp | qPCR |
| *OsJAZ8* | LOC_Os09g26780 | GAAGGCTCAACAGCTGACCAT | TTGGTGGACGGGAAGTTCTC | 69 bp | qPCR |
| *OsMYC2* | LOC_Os10g42430 | CTAGCGAGGAAACCCAATCG | CCATCCATCCATCCTAACAC | 158 bp | qPCR |
| *OsPR5* | LOC_Os12g43380 | CTGGCGGAGTTCACCATC | GCAGGAGAAGCTCATGGC | 87 bp | qPCR |
| *OsPR10* | LOC_Os03g18850 | CGGACGCTTACAACTAAATCG | AAACAAAACCATTCTCCGACAG | 174 bp | qPCR |
| *OsWRKY71* | LOC_Os02g08440 | CGGCAAAAGACGCTTGTAAC | TCGGGACCGAAGCAAATTTG | 106 bp | qPCR |
| *OsEXPB7* | LOC_Os03g01270 | GTCAGTATCCAGGGCTGACG | CGTACTCCACCAGTATCGCC | 81 bp | qPCR |
| *EXP* | LOC_Os06g11070 | TCCATCTGCTCCCGTTGTTGTG | AAAGAGTTCGCCACCAACCGTC | 273 bp | qPCR |
| *OsCOI1a* | target 1 | TGGAAGAGTGCCATATTACTG | TCAAACACAATTAATCAATGC | 496 bp | identify mutations |
| *OsCOI1b* | target 2 | TGTCATACTTTAGCTCACTGG | ACGTAAGTCCTAAGGAGCACA | 477 bp | identify mutations |
| *OsCOI2* | targets 1 and 2 | CCATGGCGTACTCCACCACG | GAGAGAACTCCTAAGACGTAG | 462 bp | identify mutations |
| *OsCOI2* | targets 3 and 4 | CCATATGTTGTCATTTGCAG | CAGCGCTGGACCAGCTGACAG | 485 bp | identify mutations |
| *Cas9* | T-DNA | GGATGATGGCATATGCAGCAG | GAGTGTGAGGTCCTGGTGGT | 1216 bp | check T-DNA insertion |
| *HPT* | T-DNA | CTCGGAGGGCGAAGAATCTC | GCTCCAGTCAATGACCGCTG | 759 bp | check T-DNA insertion |

**Table S1. List of primers used for qPCR and plant genotyping.**
